# Prion-induced heritable states in *Caenorhabditis elegans*

**DOI:** 10.64898/2026.09.27.753494

**Authors:** Orkan Ilbay, Rebecca A. Zabinsky, Thomas Lozanoski, Daniel F. Jarosz, Andrew Fire

**Affiliations:** Department of Pathology, Stanford University School of Medicine, Stanford, United States; Department of Chemical & Systems Biology, Stanford University School of Medicine, Stanford, United States; Department of Developmental Biology, Stanford University School of Medicine, Stanford, United States; Department of Genetics, Stanford University School of Medicine, Stanford, United States

## Abstract

Self-replicating protein conformations can encode persistent epigenetic states that underlie prion diseases and contribute to age-related neurodegenerative disorders in mammals. In fungi, such conformers are transmitted through mitosis and meiosis, enabling robust inheritance of diverse phenotypes. Whether analogous transgenerational protein-based inheritance extends beyond unicellular systems to metazoans remains unknown. Here, we address this question in *C. elegans*, a genetically tractable, short-lived animal well suited for tracking the transmission of heritable information across generations. We found that a yeast prion domain, when fused to a host protein, induced a heritable altered state that arose at elevated temperatures and persisted across generations at lower temperatures. The emergence of this state depended on known prionogenic yeast repeats, while the state itself was reversible and behaved semi-dominantly in crosses, consistent with a protein-based epigenetic mechanism. These findings provide direct evidence that protein domains capable of adopting self-replicating conformers can encode heritable information in an animal.

## Introduction

Epigenetic mechanisms enable genetically identical cells to adopt diverse phenotypes through stable differences in gene expression, a fundamental requirement for multicellular life (Goldberg et al., 2007). Most well-characterized epigenetic information carriers include DNA methylation, histone modifications, and small RNAs (Bošković and Rando, 2018). In addition, self-replicating protein conformations can encode heritable epigenetic states (Halfmann and Lindquist, 2010; Harvey et al., 2018; Shorter and Lindquist, 2005). These protein-based inheritance mechanisms can be remarkably stable and wide-ranging in their effects (Harvey et al., 2018; True et al., 2004), yet their scope outside fungi remains unclear (Bošković and Rando, 2018).

In the baker’s yeast *S. cerevisiae*, self-replicating protein conformers can arise spontaneously, likely during stochastic exploration of the conformational landscapes by intrinsically disordered protein domains. Once established, these conformers template their structure onto newly synthesized molecules of the same protein, triggering a self-propagating chain reaction that spreads through the cytoplasm. Yeast prions can thus be stably maintained through mitosis and meiosis, enabling robust transgenerational inheritance of diverse phenotypes, some of which can be beneficial under specific environmental conditions (Halfmann et al., 2012; True and Lindquist, 2000).

Sequence features and characteristics of prions learned from the first yeast prions (Patino et al., 1996; Wickner, 1994) have enabled the identification of a growing number of proteins capable of forming protein-based epigenetic states (Alberti et al., 2009; Chakrabortee et al., 2016). These findings suggest that dozens of yeast proteins can access self-replicating conformations, together constitute a potent layer of epigenetic regulation across diverse cellular processes (Chakravarty et al., 2020; Du et al., 2015; Garcia et al., 2021; Halfmann et al., 2012; Harvey et al., 2020; Holmes et al., 2013; Jarosz et al., 2014; Nizhnikov et al., 2016; Suzuki et al., 2012; Volkov et al., 2002).

In humans and other mammals, self-replicating protein conformations are recognized for their ability to spread between cells, and in the case of infectious prions, to transmit between individuals (Scheckel and Aguzzi, 2018), as well as for their roles in devastating neurodegenerative disorders, including prion diseases and age-related conditions such as Alzheimer’s and Parkinson’s (Jucker and Walker, 2013; Wilson et al., 2023). Moreover, animal prion-like domains can behave as bona fide prions in yeast (Si et al., 2003) and prion-like domains can support long-lived altered activity states in post-mitotic animal cells such as neurons (Khan et al., 2015). Together, these and other related studies show that animal prion-like domains are, in principle, competent to encode conformation-based persistent information and that animal cells can support the emergence and maintenance of self-replicating protein conformations.

The extent to which inheritance based on self-replicating protein conformations (also referred to as *protein-based inheritance*) operates across the tree of life remains unclear. In particular, whether such protein-based information can be transmitted through the germline and inherited across generations in an animal has not been established.

To test whether protein-based inheritance can operate in an animal, we developed an experimental framework in the nematode *C. elegans*, a genetically tractable animal with a short life cycle and invariant cell lineage, well suited for tracking the transmission of heritable information both during development and across generations (Bošković and Rando, 2018; Fitz-James and Cavalli, 2022; Heard and Martienssen, 2014; Perez and Lehner, 2019; Santilli and Boskovic, 2023).

We found that fusion to a well-characterized yeast prion domain can confer prion-like behavior on a *C. elegans* protein. The NM domain of yeast Sup35 interfered with host protein function in a temperature-dependent manner: efficient prion interference was observed at 25°C, but not at lower temperatures. Once established at 25°C, however, the altered host-protein activity state could be maintained and inherited for many generations (at least 20) at 20°C. This state depended on key prionogenic repeats in the NM domain, exhibited incomplete penetrance and ready reversibility across generations, and behaved semi-dominantly in genetic crosses. Together, these results provide direct evidence that protein domains capable of adopting self-replicating conformers can encode heritable information in an animal.

## Results

### Experimental framework to assess and study protein-based inheritance can operate in an animal model

We set out to test whether a yeast prion domain could convert a *C. elegans* protein into a prion-like element with these defining properties: (1) *de novo* emergence of a state characterized by altered protein activity,(2) dependence on key prionogenic repeats in the transplanted yeast protein, (3) inheritance of the altered protein state, (4) reversibility to the naïve state, and (5) dominance in genetic crosses. Prions could conceivably be inherited through either nuclear or extranuclear mechanisms, and we thus complete our analysis with tests that partially distinguish these inheritance modes. (**Figure 1**).

**Figure 1.**
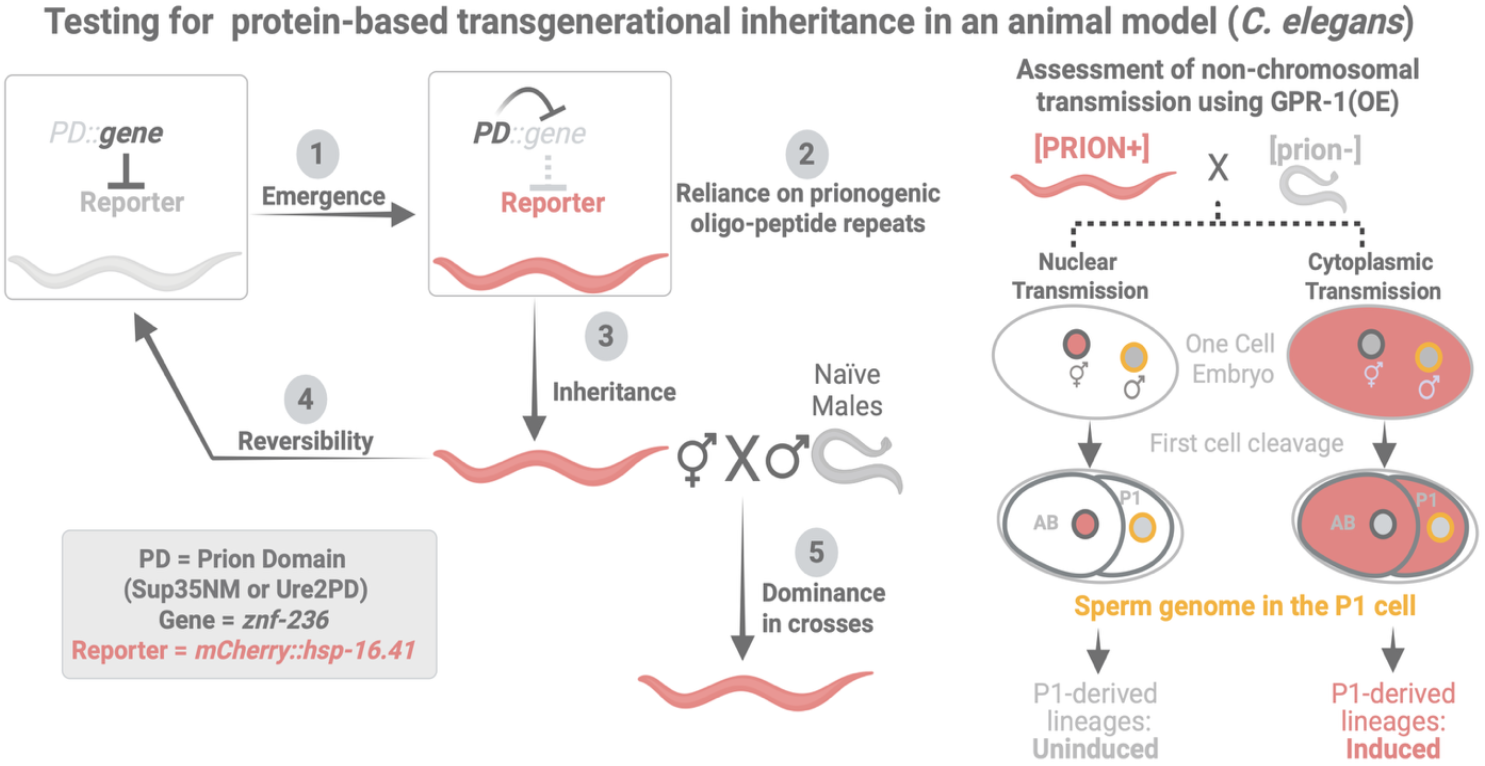
Experimental framework to assess and study protein-based inheritance in an animal model. Five defining properties of prion-like, protein-based inheritance were tested in *C. elegans*: (1) de novo emergence of an altered protein-activity state, (2) dependence on key prionogenic repeats in the transplanted yeast protein, (3) maintenance and inheritance of that state across generations, (4) reversibility to the naïve state, (5) dominance in genetic crosses. A separate outline for assaying the non-chromosomal transmission potential of the state-specifying information. A yeast prion domain (PD; Sup35NM or Ure2PD) was fused to the zinc-finger protein ZNF-236, whose activity is monitored using an endogenously tagged reporter (*mCherry::hsp-16*.*41*) that becomes de-repressed upon loss of ZNF-236 activity. In the naïve state, PD::ZNF-236 represses the reporter; spontaneous conversion to a state with reporter expression represents emergence (1). This conversion depends on the prionogenic oligo-peptide repeats within the yeast PD, tested by comparing full-length versus repeat-truncated PD fusions, representing reliance on prionogenic repeats (2). Stable propagation of the induced (reporter-expressing) state to progeny represents inheritance (3). Return of descendants to the naïve, reporter-silenced state represents reversibility (4). Crossing [PRION+] hermaphrodites to naïve males and scoring reporter induction in cross progeny tests for dominance (5). Cytoplasmic transmission was assessed by overexpressing GPR-1 [GPR-1(OE)] to force asymmetric partitioning of the sperm-derived nuclear genome into the P1 blastomere at the first embryonic cleavage, thereby uncoupling nuclear from cytoplasmic inheritance; reporter induction in P1-derived intestinal lineages indicates cytoplasmic transmission of the altered state, whereas its absence indicates nuclear (chromosomal) transmission.

To determine whether established yeast prion domains, such as the NM domain of Sup35 (Baudin-Baillieu et al., 2014) or the prion domain (N-terminal 65 amino acids) of Ure2, can interfere with protein activity in *C. elegans*, we selected the zinc-finger protein ZNF-236 as the host protein, encoded by a gene with several features that facilitate the subsequent analysis (Ilbay et al., 2026). This choice was nontrivial in that an effective host protein for such analysis must satisfy several requirements, including viability and a loss-of-function phenotype that is robust and straightforward to score.

ZNF-236 is particularly well suited for this framework because its activity can be monitored using a sensitive downstream reporter that produces readily observable phenotypes across a spectrum of ZNF-236 perturbations, from weak to strong loss-of-function alleles (Ilbay et al., 2026). Specifically, an endogenously tagged *hsp-16*.*41* reporter (*mCherry::hsp-16*.*41*) enables detection of individual worms with reduced ZNF-236 activity, which can then be isolated and propagated to test for heritability. Further, the activity of ZNF-236 is cell autonomous, allowing precise analysis of cellular function across cells and tissues.

### A yeast prion domain interferes with the activity of a host protein in *C. elegans*

The prion domains of Ure2 and Sup35, the first two yeast prions described (Patino et al., 1996; Wickner, 1994), occasionally adopt self-replicating conformations that interfere with the function of their respective proteins (**Figure 2A, B**). These prion domains can also confer prion-like behavior on heterologous functional domains when genetically fused to such domains in yeast (**Figure 2C**) (Li and Lindquist, 2000). We therefore asked whether a yeast prion domain could similarly interfere with protein function in *C. elegans* (**Figure 2D**).

**Figure 2.**
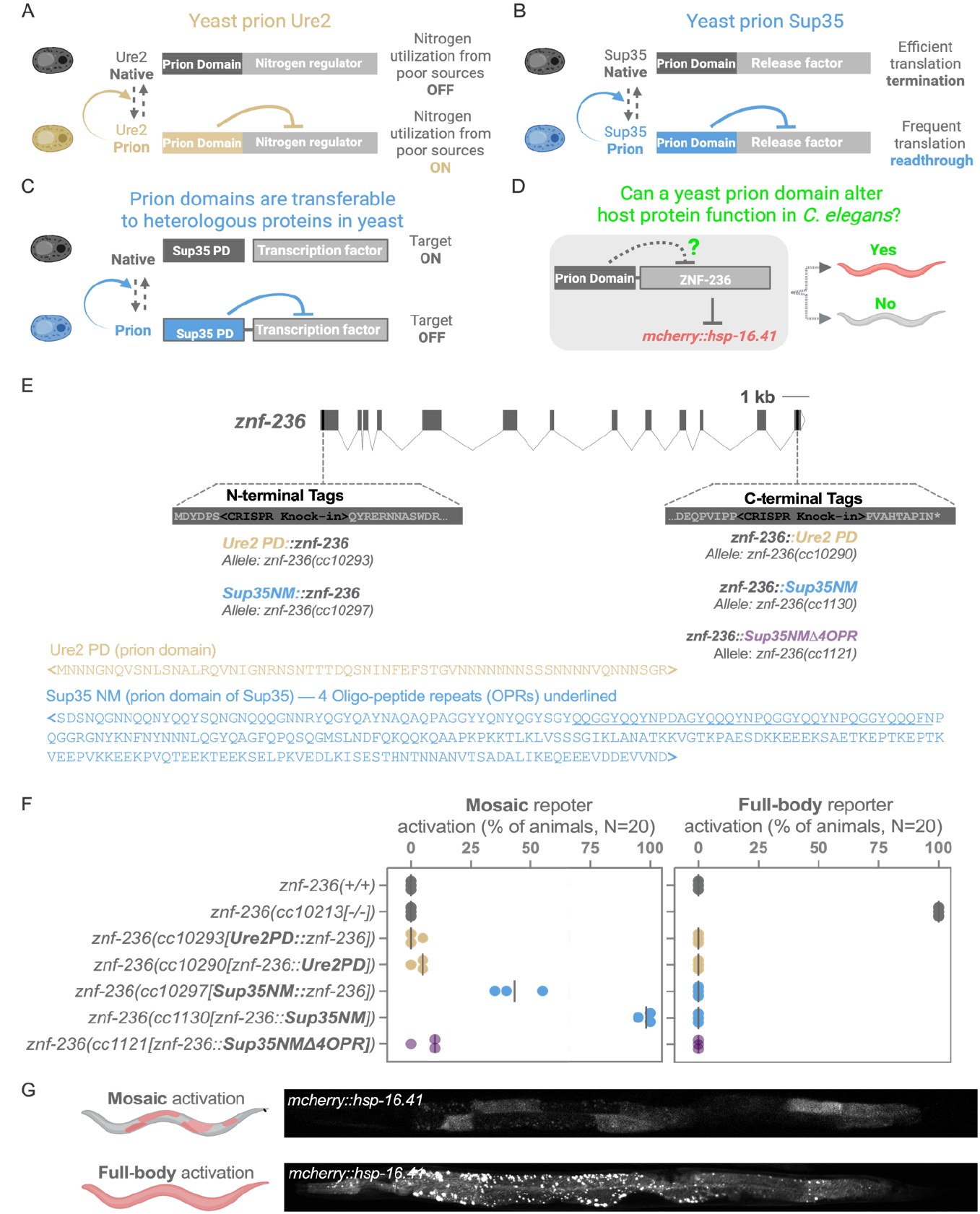
A yeast prion domain interferes with the activity of a host protein in *C. elegans*. (A–B) Yeast prion proteins as precedents for protein-based, heritable conformational switching. (A) Ure2 regulates nitrogen catabolite repression via its C-terminal nitrogen-regulator domain. In the native state, nitrogen utilization from poor sources is OFF; conversion of the N-terminal prion domain (PD) to the prion conformation derepresses the nitrogen regulator domain, switching nitrogen utilization from poor sources ON. (B) Sup35 regulates translation termination via its C-terminal release-factor domain. In the native state, translation termination is efficient; conversion to the prion conformation sequesters the release factor domain, causing frequent translational readthrough. (C) Prion domains are transferable to heterologous proteins in yeast. The Sup35 prion domain (PD), fused to an unrelated transcription factor, permits target gene activation (Target ON) in the native state; conversion of the fused PD to the prion conformation represses the transcription factor and turns the target OFF, demonstrating that a prion domain can confer prion-like, conformation-dependent regulatory switching onto an unrelated protein. (D) Experimental question: can a yeast prion domain alter host protein function in *C. elegans*? A yeast PD was fused to ZNF-236, which normally represses the reporter *mcherry::hsp-16*.*41*. If the PD confers prion-like behavior on ZNF-236 in vivo, reporter repression should be lost (Yes); if not, repression should be maintained (No). (E) Genomic structure of *znf-236* (exons, gray boxes; scale bar, 1 kb) and CRISPR knock-in strategy. PD sequences (Ure2 PD or Sup35NM) were inserted at either the N- or C-terminus of endogenous *znf-236*, generating the alleles indicated: *znf-236(cc10293)* [*Ure2PD::znf-236*, N-terminal], *znf-236(cc10297)* [*Sup35NM::znf-236*, N-terminal], *znf-236(cc10290)*[znf-236::Ure2PD, C-terminal], *znf-236(cc1130)*[znf-236::Sup35NM, C-terminal], and *znf-236(cc1121)* [znf-236::Sup35NMΔ4OPR, C-terminal], the latter lacking four oligopeptide repeats (OPRs, underlined) within the Sup35NM domain. Amino acid sequences of the Ure2 PD and Sup35NM domains used are shown, with the four OPRs deleted in the Δ4OPR allele underlined. (F) Quantification of reporter activation in each genotype at 23.6°C, scored as mosaic (partial-body) or full-body activation of *mcherry::hsp-16*.*41* (percent of animals; N = 20 per genotype). Each dot represents an independent trial, with the mean indicated by a line. *znf-236(+/+)* and *znf-236(cc10213[-/-])* serve as negative and positive controls for reporter derepression, respectively. Fusion of Sup35NM, but not Ure2PD, to ZNF-236 drives reporter activation; deletion of the four Sup35 OPRs (Sup35NMΔ4OPR) substantially reduces this activation. (G) Representative fluorescence images illustrating mosaic [top; *znf-236(cc1130)*] versus full-body [bottom; partial deletion allele, *znf-236(cc10213)]* activation phenotypes of the *mcherry::hsp-16*.*41* reporter, corresponding to the two activation categories quantified in (F).

Using CRISPR/Cas9 genome editing, we tagged *znf-236* at its endogenous locus by fusing either the Ure2 or Sup35 prion domains to the N- and C terminus of ZNF-236 (**Figure 2E**). We found that the Sup35 prion domain, but not the Ure2 prion domain, activated the reporter gene (**Figure 2F, G**). The Sup35 effect was robust to position of the tag, with both N and C terminal fusions leading to compromised ZNF-236 activity. Of the two ZNF-236 sites, the C-terminal fusion of the Sup35 prion domain produced a more penetrant interference phenotype (**Figure 2E, F**).

Unlike the full-body reporter activation observed in *znf-236* loss-of-function mutants (**Figure 2F, G**), activation in prion-tagged ZNF-236 animals was predominantly restricted to a subset of intestinal cells, resulting in a mosaic pattern. This mosaicism is consistent with cell-autonomous regulation of *mCherry::hsp-16*.*41* by ZNF-236 (Ilbay et al., 2026) and with the reporter activation only in cells in which the prion tag reduces ZNF-236 activity below critical thresholds.

Because the Sup35 prion domain’s oligopeptide repeat tract (**Figure 2E**) is markedly required for the *de novo* appearance of the [*PSI*+] state in yeast (Liu and Lindquist, 1999), we introduced the corresponding oligopeptide-repeat mutation into our *znf-236::Sup35NM* tag (**Figure 2E**) to test whether these repeats similarly modulates the frequency of reporter activation in an animal context. We note that the oligopeptide repeats (38 amino acids, 158nt with an intron) are only a fraction of the Sup35NM domain (252 amino acids, 922nt with 3 introns), so that the deletion retains the majority of the original fusion. We saw that deleting oligopeptide repeats resulted in a substantial reduction in the frequency of mosaic reporter activation (**Figure 2F**). This illustrates that the oligopeptide repeat region of the prion domain is required for efficient mosaic activation of the reporter, similar to its role in the emergence of [PSI+] in yeast (Liu and Lindquist, 1999).

### The emergence of a prion-induced state that is both heritable and reversible

The NM domain was capable of effective ZNF-236 interference in a temperature-dependent manner (**Figure 3A**). While no interference was evident at 20°C (inferred from lack of *mCherry::hsp-16*.*41* induction), and while mosaic reporter activation was prevalent at 23.6°C (**Figure 2E, F)**, we observed prominent reporter activation throughout the gut, along with activation in the excretory system at 25°C (**Figure 3A**).

**Figure 3.**
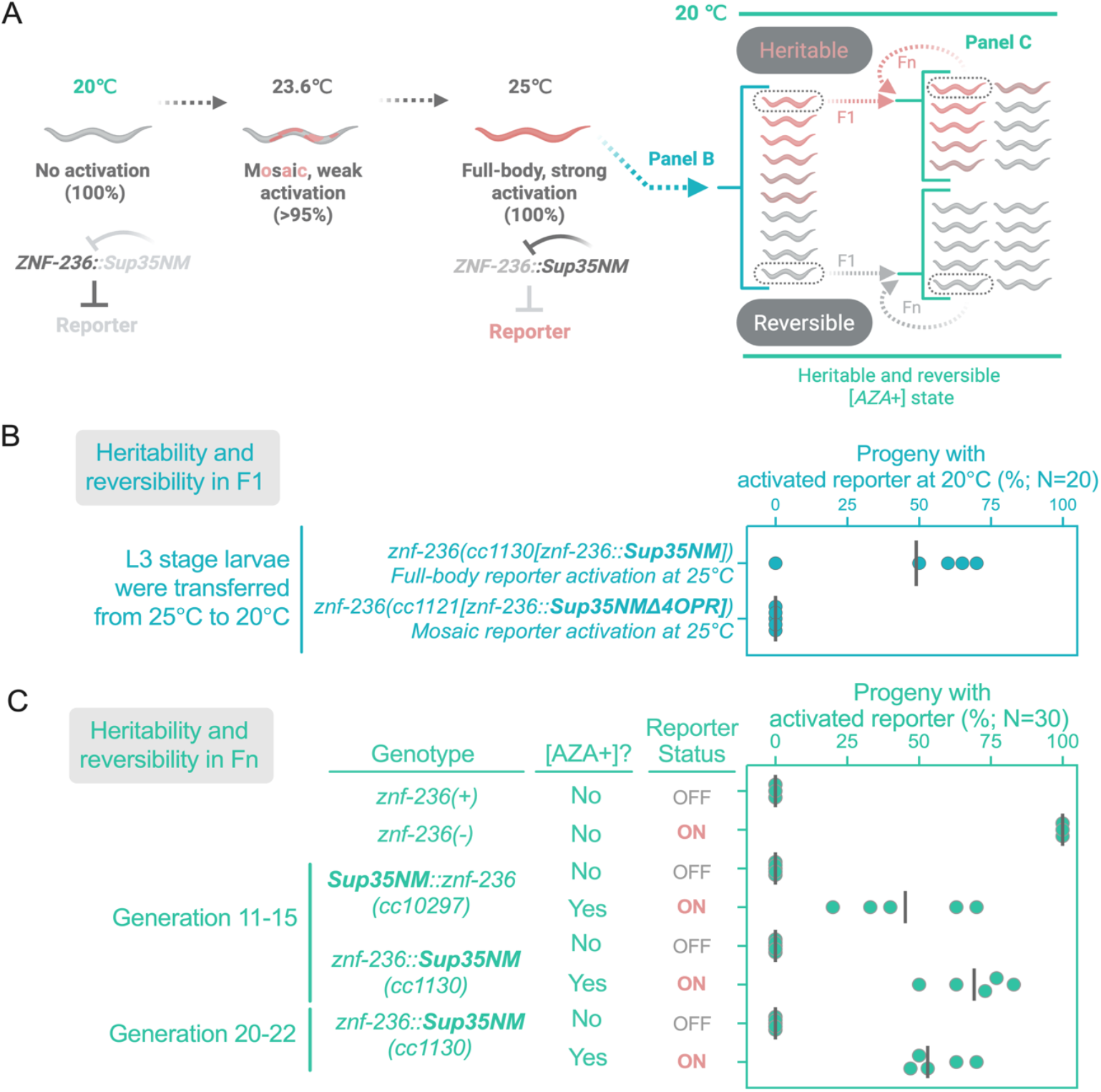
The emergence of a prion-induced state that is both heritable and reversible. **(A)** The Sup35NM domain interferes with ZNF-236 function in a temperature-dependent manner. At 20°C, ZNF-236::Sup35NM represses the *mcherry::hsp-16*.*41* reporter, and no animals showed activation (100%). At 23.6°C, mosaic, weak reporter activation predominated (>95%; see also Figure 2E, F), and at 25°C, animals showed fully penetrant, strong full-body reporter activation (100%), hereafter [*AZA*^+^] (Altered ZNF-236 Activity). Worms with the [*AZA*^+^] phenotype at 25°C were singled and transferred to 20°C, and reporter status was scored in their progeny in the F1 generation (Panel B) and in subsequent generations (Panel C). **(B)** The [*AZA*^+^] state is heritable and reversible in the F1 generation. L3-stage larvae with full-body [*AZA*^+^] activation at 25°C were transferred to 20°C, and reporter activation was scored in their F1 progeny at 20°C. Each dot represents the percentage of progeny (N = 20) scored as reporter-activated from a single parental worm; vertical lines indicate the mean. [*AZA*^+^] animals transmitted the activated-reporter phenotype to their progeny, demonstrating heritability; the state is also reversible, as [*AZA*^+^] animals gave rise to both [*AZA*^+^] and [*aza*^−^] progeny. Two genotypes were tested: *znf-236(cc1130[znf-236::Sup35NM])*, which shows full-body reporter activation at 25°C, and *znf-236(cc1121[znf-236::Sup35NMΔ4OPR])*, which shows mosaic reporter activation at 25°C. **(C)** The [*AZA*^+^] state persists across generations with variable penetrance in individual clones. Independent [*AZA*^+^] and [*aza*^−^] lineages of *Sup35NM::znf-236(cc10297)* and *znf-236::Sup35NM(cc1130)* were cultured continuously at 20°C for several generations (Fn, n = 11–15), alongside *znf-236(+)* and *znf-236(-)* controls, and reporter status was scored at generation Fn. To test whether the [*AZA*^+^] state remains heritable over more extended propagation, an independent [*AZA*^+^] lineage of *znf-236::Sup35NM (cc1130)* was additionally cultured to generations 20–22 and scored in parallel. Corresponding dot plots at right show, for each line, the percentage of progeny (N = 30) scored as reporter-activated at generation Fn. Each dot represents an independent parental worm, with horizontal lines indicating the mean.

When worms with full-body activation (hereafter [*AZA*^+^], for Altered ZNF-236 Activity) at 25°C were singled and transferred to 20°C, they transmitted the activated-reporter phenotype to their progeny (**Figure 3A, B**). Thus, the [*AZA*^+^] state was heritable. The [*AZA*^+^] state was also readily reversible: [*AZA*^+^] animals gave rise to both [*AZA*^+^] and [*aza*^−^] progeny (**Figure 3A, B**). In subsequent generations, the [*AZA*^+^] state continued to persist (at least 20 generations) with variable penetrance in individual clones (**Figure 3C**). By contrast, the mosaic activation at 23.6°C was not heritable.

### Differentially expressed genes in [*AZA*^+^] versus [*aza*^−^] states

Next, we compared gene expression between [*AZA*^+^] and [*aza*^−^] progeny of [*AZA*^+^] worms (**Figure 4**). Our goals were to determine whether the differences in *znf-236* mRNA abundance could explain the distinct induction of in *hsp-16*.*41*in [*AZA*^+^] and [*aza*^−^] progeny, whether genes induced in [*AZA*^+^] resembled those previously shown to be upregulated in *znf-236*-null mutants, and whether additional genes were differentially expressed between the two states.

**Figure 4.**
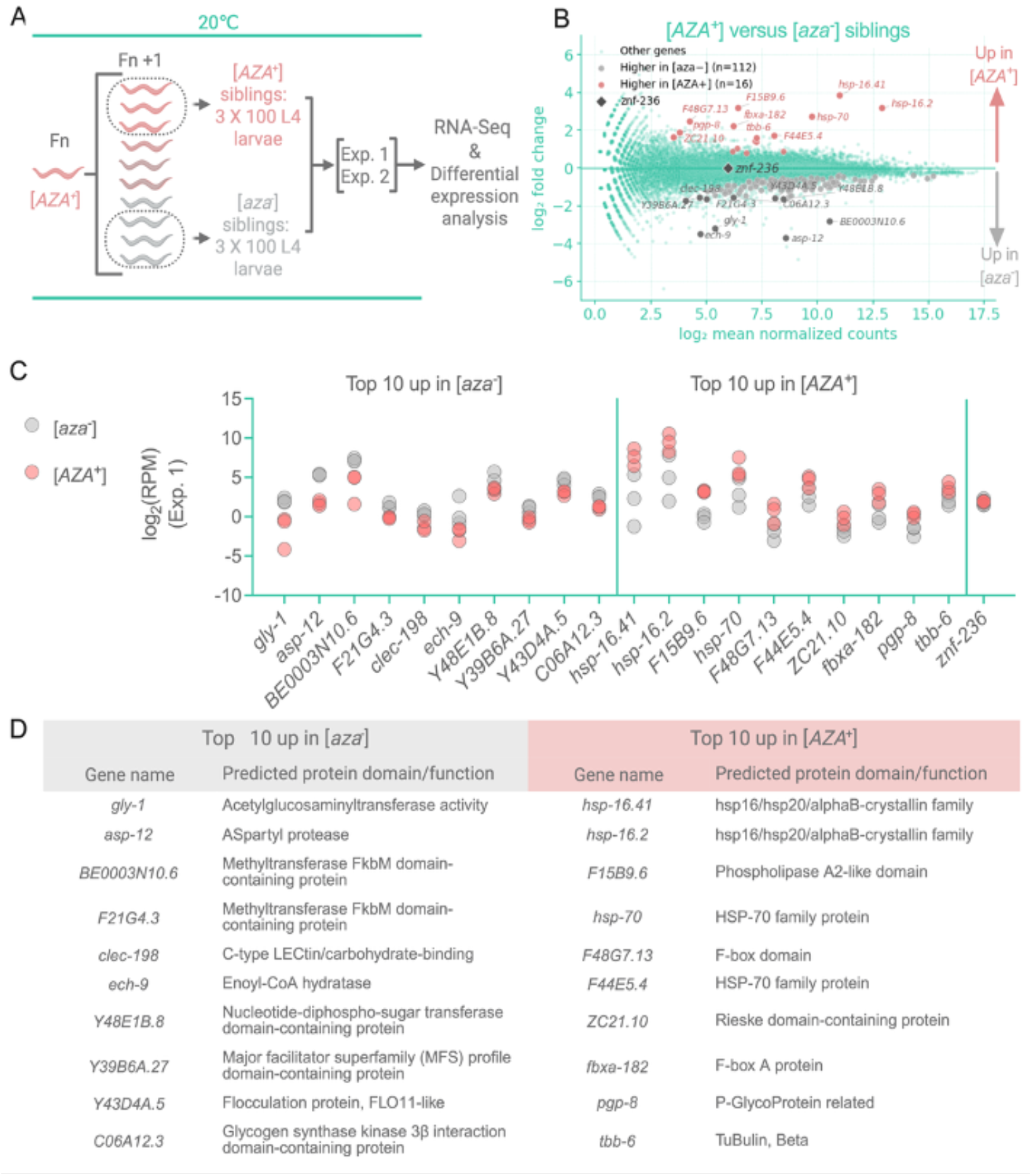
Differentially expressed genes in [*AZA*^+^] versus [*aza*^−^] states. **(A)** Experimental scheme for RNA-seq sample collection. An [*AZA*^+^] animal at generation Fn was allowed to produce progeny (Fn+1) at 20°C, which segregated into [*AZA*^+^] and [*aza*^−^] siblings. Three independent replicates of 100 L4-stage larvae each were collected separately for [*AZA*^+^] and [*aza*^−^] siblings and processed for RNA sequencing. This experiment was performed twice, independently (Exp. 1 and Exp. 2; each in triplicate); results from Exp. 1 are shown in panels B–D, and results from Exp. 2 are shown in Supplemental Figure 1. **(B)** MA plot comparing gene expression between [*AZA*^+^] and [*aza*^−^] siblings (Exp. 1). Each point represents a gene, plotted by log2 mean normalized counts (x-axis) versus log2 fold change (y-axis); positive values indicate higher expression in [*AZA*^+^] animals. Genes are colored by category: other genes (light green); genes with reproducibly higher expression in [*aza*^−^] across both experiments (gray, n = 112); genes with reproducibly higher expression in [*AZA*^+^] across both experiments (red, n = 16); and *znf-236* (black diamond). The 10 genes most strongly upregulated in [*aza*^−^] and the 10 genes most strongly upregulated in [*AZA*^+^], ranked by fold change in Exp.1, are labeled on the plot, along with *znf-236*. [*AZA*^+^]-upregulated genes include the reporter gene *hsp-16*.*41* as well as *hsp-16*.2 and hsp-70, reminiscent of the previously reported *hsp* profile in *znf-236* null worms. *znf-236* itself showed no appreciable difference in expression between [*AZA*^+^] and [*aza*^−^] siblings. **(C)** Observed expression levels, in log2(RPM), of the 10 genes most strongly upregulated in [*aza*^−^] siblings (left) and the 10 genes most strongly upregulated in [*AZA*^+^] siblings (right), ranked by fold change in Exp.1, together with *znf-236* for reference. Each dot represents an independent biological replicate; gray dots indicate [*aza*^−^] samples and pink dots indicate [*AZA*^+^] samples. Consistent with the MA plot in (B), *znf-236* expression did not differ appreciably between [*AZA*^+^] and [*aza*^−^] siblings, whereas two *hsp-16* class genes and *hsp-70* family members show elevated expression in [*AZA*^+^] animals. **(D)** Gene names and predicted protein domains/functions for the top 10 genes upregulated in [*aza*^−^] siblings (left) and the top 10 genes upregulated in [*AZA*^+^] siblings (right) shown in (C).

**Supplemental Figure 1.**
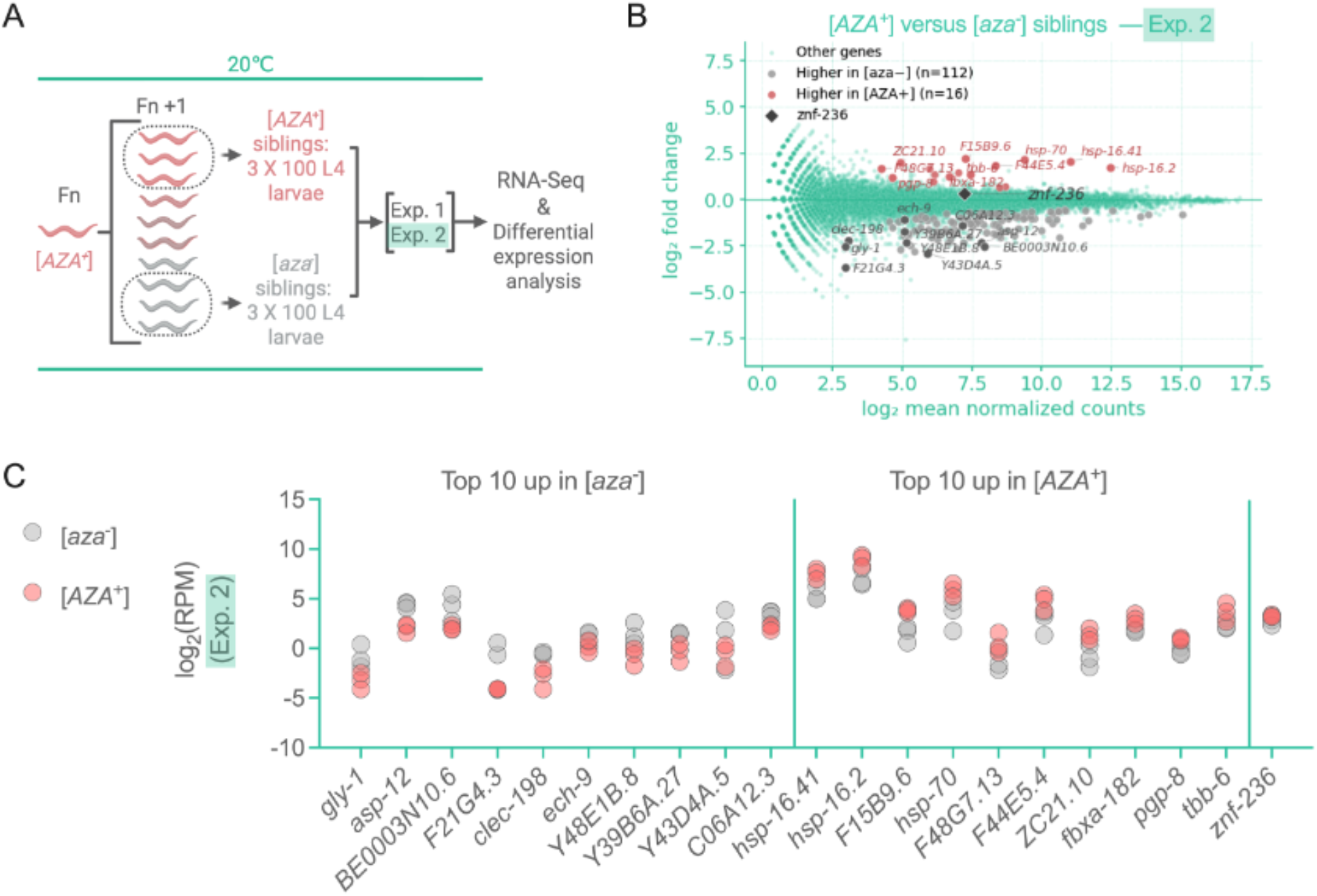
Differentially expressed genes in [*AZA*^+^] versus [*aza*^−^] siblings, independent replicate experiment (Experiment 2) **(A)** Experimental scheme for RNA-seq sample collection, as in Figure 4A. An [*AZA*^+^] animal at generation Fn was allowed to produce progeny (Fn+1) at 20°C, which segregated into [*AZA*^+^] and [*aza*^−^] siblings. Three independent replicates of 100 L4-stage larvae each were collected separately for [*AZA*^+^] and [*aza*^−^] siblings and processed for RNA sequencing, as an independent repeat (Exp. 2) of the experiment shown in Figure 4A (Exp. 1). **(B)** MA plot comparing gene expression between [*AZA*^+^] and [*aza*^−^] siblings in Experiment 2. Each point represents a gene, plotted by log2 mean normalized counts (x-axis) versus log2 fold change (y-axis); positive values indicate higher expression in [*AZA*^+^] animals. Genes are colored by category: other genes (light green); genes with reproducibly higher expression in [*aza*^−^] across both experiments (gray, n = 112); genes with reproducibly higher expression in [*AZA*^+^] across both experiments (red, n = 16); and *znf-236* (black diamond). The same 10 genes most strongly upregulated in [*aza*^−^] and 10 genes most strongly upregulated in [*AZA*^+^] identified in Experiment 1 (Figure 4B–D) are labeled here, along with *znf-236*, illustrating the concordance of expression changes between the two independent experiments. As in Experiment 1, [*AZA*^+^]-upregulated genes include the reporter gene *hsp-16*.*41* as well as *hsp-16*.*2* and *hsp-70. znf-236* itself shows no appreciable difference in expression between [*AZA*^+^] and [*aza*^−^] siblings. **(C)** Observed expression levels, in log2(RPM), of the same 10 genes most strongly upregulated in [*aza*^−^] siblings (left) and 10 genes most strongly upregulated in [*AZA*^+^] siblings (right) shown in Figure 4C, together with *znf-236* for reference, here plotted using data from Experiment 2. Each dot represents an independent biological replicate; gray dots indicate [*aza*^−^] samples and pink dots indicate [*AZA*^+^] samples. Expression patterns closely mirror those observed in Experiment 1 (Figure 4C), confirming the reproducibility of thes e gene expression changes across independent experiments

To address these questions, we collected L4-stage [*AZA*^+^] and [*aza*^−^] siblings (**Figure 4A**), extracted total RNA, and performed high-throughput RNA sequencing. To increase the statistical power and confidence in the identification of differentially expressed genes, we performed two independent experiments (Exp. 1 and Exp. 2), each comprising three biological replicates per state. Results from Exp. 1 are shown in **Figure 4**, and results from Exp. 2 are shown in **Supplemental Figure 1**. We analyzed each experiment independently using DESeq2 and defined high-confidence differentially expressed genes as those with a nominal *P* value < 0.05 in in each experiment, a concordant direction of change, and a Fisher-combined *P* value that passed Benjamini–Hochberg correction at an FDR of 0.05. This analysis identified 128 reproducibly differentially expressed genes in [*AZA*^+^] relative to [*aza*^−^] siblings, of which 16 were upregulated and 112 were downregulated (Supplemental Data 1). Forty-one of these genes exhibited at least a twofold change in both experiments. The relative expression levels and predicted functions of the 10 genes most strongly upregulated genes in either [*aza*^−^] or [*AZA*^+^] siblings, ranked by fold change in Exp. 1, are shown in **Figure 4C** and **4D**.

RNA-Seq analysis revealed no detectable difference *znf-236* mRNA levels between [*AZA*^+^] and [*aza*^−^] worms, indicating that differences in the steady-state abundance of the *znf-236* transcript are unlikely to account for phenotypic differences between these two states (Figure 4B, 4C, and S1).

We next asked whether [*AZA*^+^] animals showed transcriptional changes consistent with reduced ZNF-236 activity. Transcript levels of several members of the *hsp-16* class chaperone family, as well as *hsp-70*, were elevated in [*AZA*^+^] animals (**Figure 4B, 4C, and S1**). This expression signature is consistent with compromised ZNF-236 activity (Ilbay et al., 2026). Expression of these *hsp* genes varied substantially among [*aza*^−^] samples, suggesting that the robustness of their default silencing in animals carrying ZNF-236::Sup35NM may be compromised, rendering their expression particularly sensitive to subtle environmental changes, including changes in temperature.

Beyond these *hsp* genes, we identified additional genes whose expression changed consistently between the two states across both experiments (**Figure 4C, D and S1**). Several of these genes, including *asp-12* and *F15B9*.*6*, have not previously been implicated as targets of ZNF-236. These gene expression changes may represent a response to [*aza*^−^]-to-[*AZA*^+^] switch or identify pathways that contribute to the maintenance or reversal of the [*AZA*^+^] state.

### The prion-induced heritable state [*AZA*^+^] acts semi-dominantly in crosses

Yeast prions act dominantly in crosses, consistent with conversion of natively folded proteins in the naive cytoplasm into self-replicating conformers. We therefore tested whether the [*AZA*^+^] state is dominant in *C. elegans*.

Notably, when *znf-236(*−*/*−) hermaphrodites were crossed with *znf-236(+/+)* males, we did not observe reporter activation in heterozygous cross progeny (**Figure 5A**). This indicates that *znf-236* is haplosufficient (**Figure 5B**), such that ZNF-236 produced from a single copy of the gene is sufficient to prevent reporter activation.

**Figure 5.**
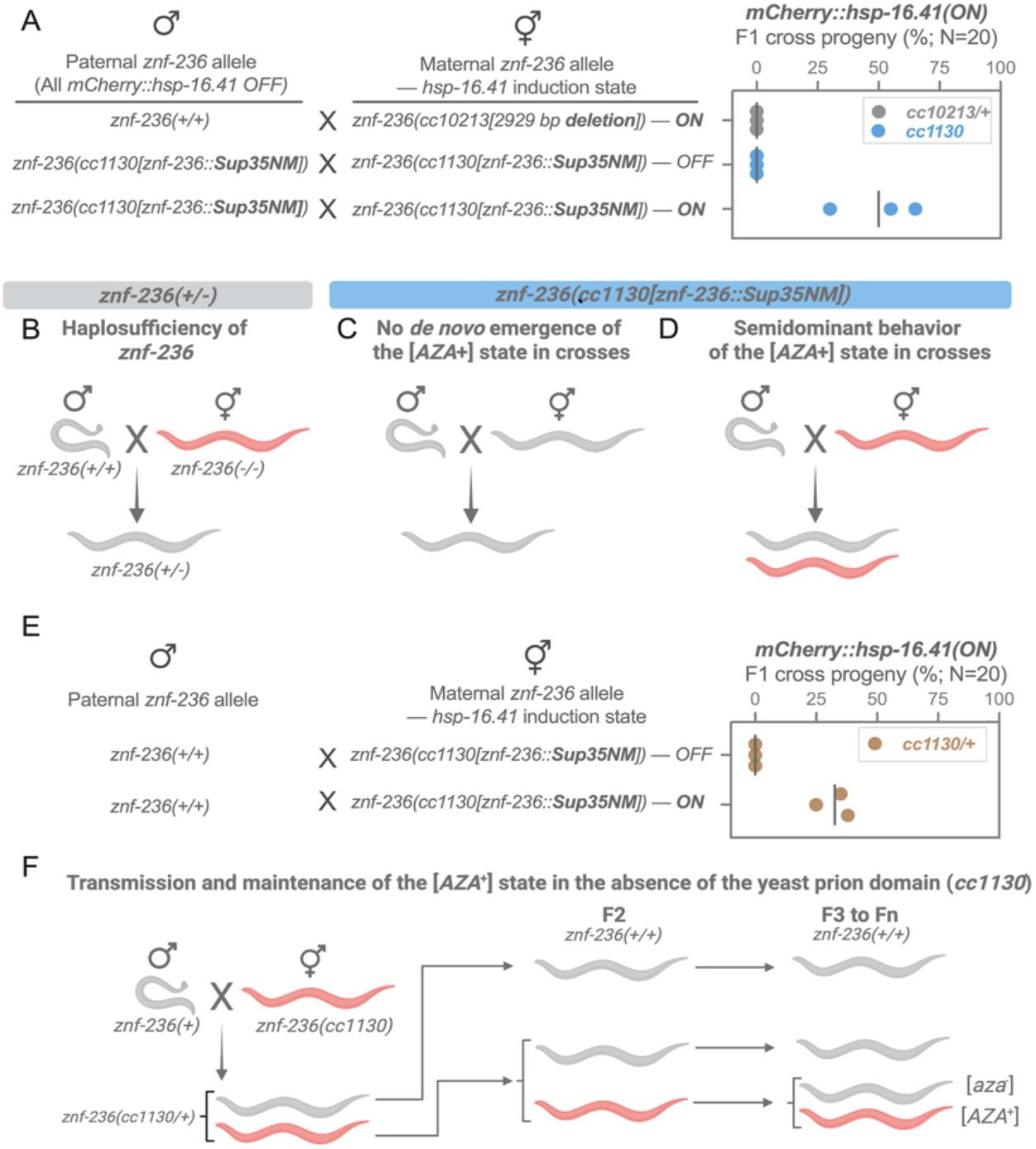
Semi-dominant transmission in genetic crosses and maintenance of the [*AZA*^+^] state in the absence of the yeast prion domain. **(A)** Reporter (*mCherry::hsp-16*.*41*) induction in F1 cross progeny from reciprocal crosses testing haplosufficiency, de novo emergence, and dominance of the [*AZA*^+^] state. Paternal *znf-236* alleles (all *mCherry::hsp-16*.*41* OFF) were crossed to hermaphrodites carrying the indicated maternal *znf-236* allele and reporter induction state (ON/OFF): *znf-236(+/+)* males × *znf-236(cc10213[2929 bp deletion])* hermaphrodites (reporter ON in the maternal strain); *znf-236(cc1130[znf-236::Sup35NM])* males × *znf-236(cc1130[znf-236::Sup35NM])* hermaphrodites in the [*aza*^−^] (reporter OFF) state; and the same cross with hermaphrodites in the [*AZA*^+^] (reporter ON) state. Percentage of F1 cross progeny with an activated reporter is shown at right (N = 20); each dot represents an independent cross, with the vertical line indicating the mean. **(B)** Haplosufficiency of *znf-236*. Crossing *znf-236(+/+)* (strain name: PD9295) males with *znf-236(-/-)* (strain name: PD10213) hermaphrodites yields *znf-236(+/-)* heterozygous progeny with no reporter activation, indicating that a single wild-type copy of *znf-236* is sufficient to repress the reporter. **(C)** No de novo emergence of the [*AZA*^+^] state in crosses. Crossing [*aza*^−^] *znf-236(cc1130[znf-236::Sup35NM])* (strain name: PD1130) hermaphrodites with [*aza*^−^] males of the same *znf-236* genotype (strain name: PD10207) yields only [*aza*^−^] progeny, indicating that mating alone does not trigger de novo emergence of the [*AZA*^+^] state. **(D)** Semidominant behavior of the [*AZA*^+^] state in crosses. Crossing [*AZA*^+^] *znf-236(cc1130[znf-236::Sup35NM])* (strain name: PD1130) hermaphrodites with [*aza*^−^] males of the same *znf-236* genotype (strain name: PD10207) yields a substantial fraction of [*AZA*^+^] progeny, indicating semi-dominant transmission of the altered ZNF-236 state. **(E)** Interference with wild-type ZNF-236 activity does not require the yeast prion domain. *znf-236(+/+)* males lacking the *cc1130* allele were crossed to *znf-236(cc1130[znf-236::Sup35NM])* hermaphrodites in either the [*aza*^−^] (OFF) or [*AZA*^+^] (ON) state, yielding *cc1130/+* heterozygous F1 progeny. A substantial fraction of F1 progeny from [*AZA*^+^] mothers displayed reporter activation, despite inheriting a paternal *znf-236* allele that encodes wild-type ZNF-236 with no Sup35NM. Percentage of F1 cross progeny with an activated reporter is shown at right (N = 20); each dot represents an independent cross, with the vertical line indicating the mean. **(F)** Transmission and maintenance of the [*AZA*^+^] state in the absence of the yeast prion domain. *znf-236(+)* males were crossed to [*AZA*^+^] *znf-236(cc1130)* hermaphrodites, yielding *cc1130/+* heterozygous F1 progeny that segregate into [*aza*^−^] and [*AZA*^+^] siblings. These F1 hermaphrodites were allowed to self, generating F2 progeny that segregate for the *znf-236* locus, including *znf-236(+/+)* homozygotes that no longer carry the *cc1130* allele or the Sup35NM prion domain. Both reporter-ON and reporter-OFF states persisted among these *znf-236(+/+)* F2 animals and were stably maintained through subsequent generations (F3 to Fn), demonstrating that once established, the [*AZA*^+^] state can be propagated independently of the yeast prion domain that was required to initiate it.

Next, we crossed hermaphrodites carrying the *cc1130* allele (ZNF-236::Sup35NM) with males of the same genotype. When *cc1130* hermaphrodites were in the [*aza*^−^] state, all cross progeny were also [*aza*^−^] (**Figure 5A, C**), indicating that mating itself does not trigger *de novo* emergence of [*AZA*^+^]. In contrast, when [*AZA*^+^] hermaphrodites were crossed to [*aza*^−^] males, a large fraction of the cross progeny were also [*AZA*^+^] (**Figure 5A, D**), indicating a semi-dominant transmission consistent with prion-like spreading of the altered ZNF-236 state.

### The [*AZA*^+^] state is maintained in the absence of the yeast prion domain

The semi-dominant inheritance of the [*AZA*^+^] state suggests that the [*AZA*^+^] state can interfere with the ZNF-236 activity when the *cc1130* allele, which encodes ZNF-236::Sup35NM, is introduced by naive males in crosses. We next asked whether Sup35NM was necessary for this cross-inhibition or whether the [*AZA*^+^] state could similarly interfere with wildtype ZNF-236 activity.

In crosses between [*AZA*^+^] hermaphrodites and males that lacked the *cc1130* allele and were wildtype for *znf-236* (**Figure 5E**), a fraction of the cross progeny also displayed the [*AZA*^+^] phenotype (**Figure 5E**). This indicates that the [*AZA*^+^]-associated interference with ZNF-236 activity does not require the yeast prion domain. We further found that the [*AZA*^+^] state was maintained in the F2 progeny that were wild-type for *znf-236* (**Figure 5F**). Thus, maintenance of the [*AZA*^+^] state at 20°C does not require Sup35NM, even though this yeast prion domain was needed for the observed induction of the state at 25°C.

One possible explanation is that Sup35NM initiates a prion-like state that can subsequently be maintained by ZNF-236 itself. ZNF-236 contains multiple intrinsically disordered regions (IDRs), and some IDRs are known to support prion-like behavior in yeast (Chakrabortee et al., 2016). Moreover, ZNF-236’s domain architecture (zinc-finger domains interspersed by IDRs) resembles that of the yeast protein, Azf1, whose transient overexpression can induce the prion-like heritable state [*AZF1*^+^] (Chakrabortee et al., 2016). These observations raise the possibility that Sup35NM acts as an initial trigger that converts ZNF-236 into a self-propagating state capable of persisting after the yeast prion domain is no longer present.

### Non-chromosomal transmission of the prion-induced [*AZA*^+^] state

Yeast prions can undergo cytoduction: in a specialized mutant background, the cytoplasms of two mating partners transiently mix while nuclear fusion is prevented, allowing prion conversion in the absence of genetic mixing (Conde and Fink, 1976; Tuite et al., 1983). In this setting, prion-negative (naïve) cells can be converted to the prion state through cytoplasmic exchange with prion-positive cells.

A functionally analogous experiment can be performed in *C. elegans* using tools that enable non-Mendelian segregation of parental genomes. Germline overexpression of *gpr-1* [GPR-1(OE)] can prevent mixing of maternal and paternal genomes after fertilization, resulting in segregation of the male and hermaphrodite genomes into separate blastomeres at the first embryonic division (Artiles et al., 2019; Besseling and Bringmann, 2016). The two resulting blastomeres (AB and P1) give rise to distinct lineages; notably, the intestine and germline descend from P1. Thus, when the P1 lineage inherits only the sperm genome from an [*aza*^−^] male, appearance of an [*AZA*^+^] phenotype in the intestine would indicate that the activated state can be transferred to a lineage that did not inherit the maternal chromosomes, consistent with non-chromosomal transmission through the ooplasm. If the transferred state also enters the P1-derived germline and is maintained through gametogenesis, F2 progeny derived from these non-Mendelian F1 animals would likewise be expected to display the [*AZA*^+^] state.

We tested these predictions using control crosses between *znf-236(-/-); GPR-1(OE)* hermaphrodites and *znf-236(+/+)* males (**Figure 6A; Supplemental Table 1**). In F1 progeny in which the P1 lineage carried only the paternal (wild-type) genome, the intestine did not display the znf-236(null) phenotype (**Figure 6A, B**). By contrast, all progeny in which P1 inherited maternal *znf-236(-/-)* genome showed reporter activation in the intestine and produced reporter-activated F2 progeny. Thus, in these control crosses using a genetic null allele, the intestine and F2 phenotypes corresponded to the parental origin and znf-236 genotype of the genome inherited by P1. These results establish the expected pattern for chromosomal transmission of the *znf-236(null)* phenotype.

**Figure 6.**
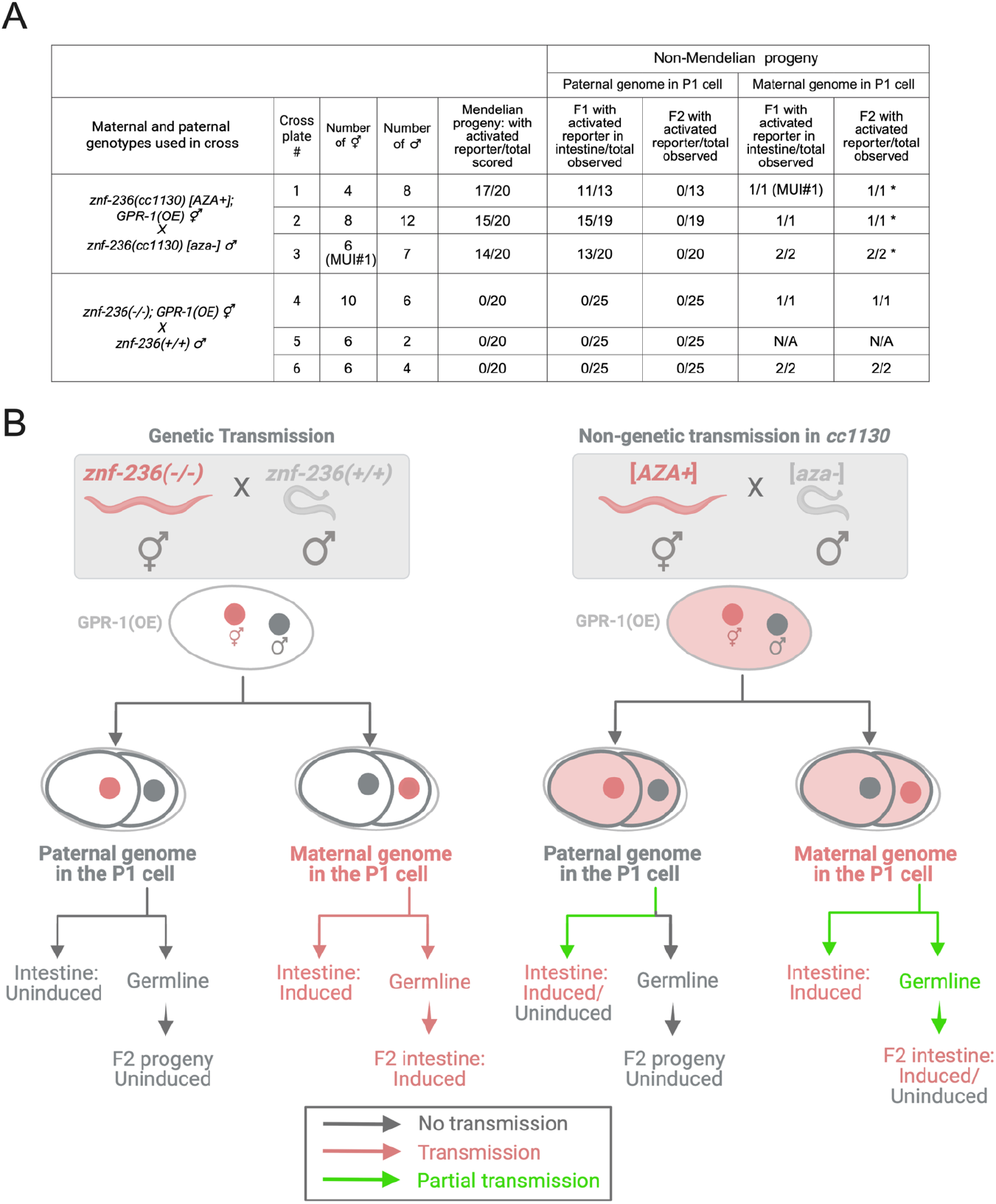
Non-chromosomal transmission of the prion-induced [*AZA*^+^] state. **(A)** Crosses testing transmission of the activated *znf-236* reporter. Cross plates 1–3: *znf-236(cc1130); GPR-1(OE)* hermaphrodites carrying the reporter in an active epigenetic state [*AZA*^+^], induced by growth at 25°C, were crossed to *znf-236(cc1130)* males carrying the reporter in the inactive state [*aza*^−^]. Cross plate 3 used a maternal uniparental isodisomy (MUI) line (MUI#1), derived from a hermaphrodite that had inherited both copies of the relevant chromosome from its mother, with no paternal contribution. Cross plates 4–6 are Mendelian control crosses between *znf-236(-/-); GPR-1(OE)* hermaphrodites and *znf-236(+/+)* males. For each cross, the number of Mendelian progeny with an activated reporter is shown, along with the fraction of non-Mendelian progeny (scored by parent-of-origin of genome in the P1 cell, as diagrammed in B) showing an activated reporter in the F1 intestine and in the F2 generation. Asterisks indicate F1 animals that produced a mixture of reporter-positive and reporter-negative F2 progeny, demonstrating incomplete transmission of the activated state and recapitulating the variable inheritance characteristics of [*AZA*^+^]. (B)Schematic summarizing reporter activation according to the parental origin of the genome present in the P1 cell. Left: In control crosses between *znf-236(−/−); GPR-1(OE)* hermaphrodites and *znf-236(+/+)* males, the maternal and paternal genomes segregate between the AB and P1 blastomeres. The paternal genome is more frequently present in the P1 lineage than the maternal genome (Artiles et al., 2019). F1 animals with the paternal genome in the P1 cell had an uninduced intestine and produced uninduced F2 progeny. F1 animals with the maternal genome in the P1 cell had an induced intestine and produced induced F2 progeny. Right: In crosses between [*AZA*^+^] hermaphrodites and [*aza*^−^] males, F1 animals with the paternal genome in the P1 cell included animals with induced and uninduced intestines; their F2 progeny were uninduced. F1 animals with the maternal genome in the P1 cell had induced intestines and produced both induced and uninduced F2 progeny. Arrow colors summarize the observed outcomes: gray, no transmission; salmon/red, transmission; green, partial or variable transmission.

Next, we crossed *znf-236(cc1130[ZNF-236::Sup35NM]); GPR-1(OE)* [*AZA*^+^] hermaphrodites with [*aza*^−^] males (**Figure 6A**). Among the Mendelian F1 progeny, the majority of the animals displayed reporter activation (Figure 6A), consistent with the semi-dominant inheritance of the maternally derived [*AZA*^+^] state. We then examined the non-Mendelian progeny in which the P1 lineage inherited the paternal genome from the [*aza*^−^] male. Despite carrying the genome of the inactive parent, 39 of 52 of these animals displayed reporter activation in the F1 intestine (Figure 6A, B). Reporter activation was observed in all three independent crosses, including the cross performed with the MUI#1 (maternal uniparental isodisomy) line. In contrast, intestinal activation was never observed when the paternal znf-236(+/+) genome was inherited in the control crosses (0/75) (**Figure 6A**). These results indicate that the activated state can pass from the [*AZA*^+^] hermaphrodite to an embryonic lineage containing the paternal genome of [*aza*^−^] male, supporting non-chromosomal transmission through the maternal ooplasm.

We next asked whether the activated state acquired by the paternal-genome P1 lineage was transmitted through its germline. Although reporter activation was frequent in the F1 intestine, none of the corresponding animals produced reporter-activated F2 progeny (0/52). Thus, under these conditions, the maternally supplied ooplasm contained activity sufficient to produce the [*AZA*^+^] phenotype in a somatic lineage containing the [*aza*^−^] paternal genome but was not detectably propagated through the germline to the next generation (Figure 6A, B).

A different outcome was observed when P1 inherited the maternal genome from [*AZA*^+^] worms (a rare configuration observed in a small fraction of the progeny from a cytoduction cross in Fig 6). All four such F1 animals examined displayed reporter activation in the intestine and produced reporter-activated F2 progeny. However, these F1 animals produced mixtures of reporter-on and reporter-off F2 progeny rather than uniformly activated populations. This incomplete transmission differed from the uniform reporter activation observed in F2 progeny carrying the maternal *znf-236* null allele. Moreover, because these non-Mendelian animals carried a uniparental isodisomic genome, the reporter-on and reporter-off phenotypes among their descendants are unlikely to result from the segregation of genetic differences or parental polymorphisms. Instead, the variable phenotype recapitulated the incomplete transmission characteristic of [*AZA*^+^], supporting inheritance of a metastable and non-genetic state rather than a conventional *znf-236* mutation.

## Discussion

In this study, we addressed whether inheritance based on self-replicating protein conformations could operate in an animal. We found that the yeast prion domain Sup35NM can interfere with the activity of the *C. elegans* protein ZNF-236, leading to reporter activation and gene-expression changes resembling those caused by partial loss-of-function alleles of *znf-236*. Efficient interference with host protein function was induced only at a higher culture temperature (25°C), yet the resulting altered state is heritable, persisting for at least 20 generations at the standard culture temperature for *C. elegans* (20°C). Several observations support a prion-like interpretation of this state, which we named [*AZA*^+^]: (i) the emergence of the [*AZA*^+^] state requires the prionogenic repeat sequences of Sup35NM, which constitute a relatively small portion (∼15%) of this prion domain. Beyond this, the [*AZA*^+^] state showed (ii) semi-dominance, (iii) reversibility, and (iv) cytoplasmic transmission, inheritance properties that are not observed in genetic loss-of-function alleles of *znf-236*. Our findings support a model in which Sup35NM promotes conversion of the ZNF-236 fusion protein from a naïve state to a self-propagating, prion-like state, analogous to the conformational conversion mediated by Sup35NM in yeast.

We used a well-studied yeast prion as a candidate trigger to test for protein-based inheritance, and we observed prion-like patterns of inheritance induced by a yeast prion domain. These results suggest that the heritable state that we report here is induced by self-replicating protein conformations acting as epigenetic information carriers, a mechanism well established in yeast. Nonetheless, such conformations have not yet been directly characterized in *C. elegans*, and we cannot formally rule out the involvement of other, non-protein-based epigenetic information carriers. Conventional epigenetic mechanisms that are known to operate in *C. elegans* include histone modifications and small RNAs, which generally act transcriptionally or post-transcriptionally to alter steady-state mRNA levels of target genes. However, *znf-236* mRNA levels are comparable between [*AZA*^+^] and [*aza*^−^] siblings, arguing against a mechanism in which the altered state is maintained primarily through changes in *znf-236* transcription of mRNA stability. This observation does not exclude indirect contributions from chromatin- or RNA-based pathways, including effects on other genes that regulate ZNF-236 protein abundance or activity.

One could also speculate that the expression of the NM domain of Sup35 alone—independent of ZNF-236, and regardless of whether the underlying mechanism is protein-based or conventional—could trigger a heritable response in the worm. To study the potential prion-like behavior and cell-to-cell transmission of the NM domain of Sup35, Nussbaum-Krammer et al. previously expressed this domain in *C. elegans* under strong tissue-specific promoters from multicopy extrachromosomal arrays (Nussbaum-Krammer et al., 2013). Under these conditions, Sup35NM formed highly toxic aggregates that spread between cells in *C. elegans*, however heritable changes in this case were not described (Nussbaum-Krammer et al., 2013).

A recent study from Eroglu *et al*. reported that endogenous amyloid-like protein structures, visible in the gonads of wild-type worms and mutants that show enhanced accumulation of such structures, can be biochemically isolated and introduced into the gonads of wild-type worms to induce specific phenotypes (Eroglu et al., 2024). These phenotypes are inherited for a few generations. This finding suggests epigenetic inheritance mediated by proteinaceous vehicles present in amyloid extracts, although whether these vehicles contain prion-like particles remains unclear. We don’t know if prion-induced heritable states we report in this study and the inheritance mediated by the amyloid-like protein structures reported by Eroglu *et al*. share any underlying fundamental principles. Nonetheless, the prion-induced states we report here appear to differ in their heritability, persisting for at least 20 generations compared to a few generations reported by Eroglu et al.

The basis for the temperature dependence of [*AZA*^+^] emergence is not yet clear. One possibility is that 25°C presents mild but sufficient stress to overwhelm certain protein-quality-control pathways that otherwise prevent the formation of self-replicating Sup35NM seeds. Alternatively, elevated temperature might directly promote the folding of Sup35NM into self-replicating conformers. Temperature could also affect the abundance, localization, turnover, or interacting partners of ZNF-236::Sup35NM, thereby altering its susceptibility to conversion. Previous studies have shown that mild heat stress (23°C-25°C), heat-shock (35°C-37°C), or other environmental stresses can establish heritable responses, including altered gene expression, in *C. elegans*. These responses can involve germline histone modification, and small-RNA pathways. For example, transient growth at 25°C can produce changes in mRNA abundance that persists several generations after animals are returned to 20°C; these effects depend on endogenous germline RNAi (Schott et al., 2014). Temperature-induced expression of extrachromosomal arrays can also persist for many generations and is associated with reduced H3K9me3 at the affected array chromatin (Klosin et al., 2017). Maternal heat stress can also promote HSF-1 and MET-2-dependent H3K9me2 bookmarking that persists in progeny (Das et al., 2021), whereas HRDE-1-dependent nuclear RNAi limits the transgenerational persistence of heat-induced transcriptional activation (Ni et al., 2016). Other stress-induced phenotypes require soma-to-germline signaling and germline chromatin or small-RNA machinery for their inheritance (Kishimoto et al., 2017; Okabe et al., 2021). Heritable small-RNA responses can display stochastic differences in persistence among genetically identical lineages, resembling the incomplete penetrance in [*AZA*^+^], albeit with variation occurring primarily between maternal lineages rather than among progeny of the same mother (Houri-Zeevi et al., 2020).

These mechanisms raise the possibility that growth at 25°C creates an inherited RNA- or chromatin-based state that facilitates the establishment or phenotypic expression of [*AZA*^+^]. Such a mechanism need not be mutually exclusive with protein-based induction or inheritance: a heat-induced epigenetic response could increase the probability that ZNF-236::Sup35NM converts to a self-replicating conformation. Alternatively, an altered ZNF-236::Sup35NM conformer could engage RNA- or chromatin-based pathways to reinforce its effects. Nonetheless, the specific requirement for the Sup35NM fusion, together with semi-dominance and non-chromosomal transmission of altered ZNF-236 activity, supports a model in which the tagged protein participates directly in initiating [*AZA*^+^], although these observations do not establish that the protein is the sole carrier of the state.

The reversibility of the [*AZA*^+^] state is high—surprisingly high compared to many prion states described in yeast. This might be due to unequal transmission of [*AZA*^+^]-specifying factors (e.g., prion-like, self-replicating protein conformations) into germ cells, frequently leaving the amount of such factors below a certain threshold required for stable propagation. This unequal transmission could be coupled with differences in the capacity of germ cells to mitigate the impact of such factors through protein quality control or other homeostatic mechanisms. A second, non-exclusive possibility is that the [*AZA*^+^] state is disease-like: an aberrant epigenetic state triggered by a prion domain under mild stress (e.g., elevated temperature). If so, it is conceivable that cellular homeostasis mechanisms actively counteract this state, and that this response underlies the frequent reversions to a naïve state.

The transmission of the [*AZA*^+^] state is evident in the intestine but not in the germline of the chimeric F1 generation (inferred from the absence of the [*AZA*^+^] in F2s), even though both tissues derive from the P1 blastomere, which carries the naïve paternal genome while inheriting the ooplasm of the [*AZA*^+^] hermaphrodite. Although transmission of the activated reporter phenotype of the [*AZA*^+^] state to the intestines of chimeric worms (**Figure 6**) demonstrates the non-chromosomal, cytoplasmic transmission potential of the [*AZA*^+^] state, the lack of transmission of the [*AZA*^+^] state to the next generation (through the germline) suggests that shared cytoplasm alone may not always be sufficient. ZNF-236 contains multiple C2H2 zinc-finger domains and is therefore predicted to associate with DNA and/or chromatin, raising the possibility that maternal chromosomes could carry chromatin-associated ZNF-236::Sup35NM in either its naïve or prion-like conformational state. Nuclear localization may limit cytoplasmic transmission, as nuclear-residing yeast prions can be transmitted less efficiently by cytoduction than predominantly cytoplasmic prions (Radchenko et al., 2011; Rogoza et al., 2010). One potential explanation is that [*AZA*^+^]-specifying factors (e.g., self-replicating conformers of the ZNF-236::Sup35NM fusion) exist in two pools (cytoplasmic and chromatin- or nucleus-associated) and both pools may be required for the transmission of the [*AZA*^+^] state to the germline of the next generation.

Alternatively, although cytoplasmic induction is sufficient to produce the activated reporter phenotype in the intestine, the transmission of the [*AZA*^+^] state to the germline of offspring may instead depend on factors associated with the inherited maternal chromosomes, which could act either independently or in combination with cytoplasmically inherited factors. In either model, cytoplasmically inherited factors may be particularly susceptible to germline-specific proteostasis mechanisms that eliminate protein aggregates during oocyte maturation (Bohnert and Kenyon, 2017), thereby potentially limiting their transmission to the F2 generation.

The *C. elegans* genome encodes hundreds of prion-like domains, disordered protein sequences that share compositional similarities with the NM domain of Sup35 and other known prion domains. We hypothesize that at least some of these domains are capable of adopting self-replicating protein conformations, granting access to prion-like epigenetic states. The ability of the yeast prion domain Sup35NM to induce heritable interference with ZNF-236 activity provides a platform to test whether endogenous *C. elegans* prion-like domains can similarly trigger heritable interference when fused to ZNF-236. Reporters that track naïve-to-prion-like conversion of candidate proteins, combined with systematic screens for triggering conditions, should help reveal how widely animals exploit this mode of inheritance. This system described here should also allow identification of genetic and epigenetic mechanisms modulating prion emergence and/or propagation (e.g., HSP104 and PIN1 respectively)(Chernoff et al., 1995; Derkatch et al., 2001).

The maintenance of the [*AZA*^+^] state in the absence of the triggering yeast prion (**Figure 5**) suggests that ZNF-236 may acquire prion-like behavior, making it a strong candidate for an endogenous *C. elegans* protein with prion-like properties. Switching to altered activity states for proteins bearing potentially functional prion-like sequences in the *C. elegans* genome could similarly be triggered by temperature stress, as well as by other environmental, developmental, or physiological inputs. This would allow worms to establish long-lasting changes in protein activity that are transmitted across generations yet remain readily reversible, enabling phenotypic diversity in response to relevant stimuli.

## Acknowledgments

We thank Victor Ambros (UMass Chan Medical School), Dae-Eun Jeong (Stanford University), Haim Krupkin (Stanford University), Alexandria Van Elgort (Stanford University), and Richard I. Morimoto (Northwestern University) for critical reading of the manuscript and helpful comments. This work was supported by the National Institutes of Health through grants R35GM130366 (A.F.) and R01AG086495 (D.J.).

## Declaration of generative AI use

During the preparation of this work, the authors used ChatGPT (GPT-5; OpenAI), under human supervision, to assist with editing portions of the manuscript and drafting Python and R scripts used in the RNA-seq analysis, including the DESeq2 workflow. The authors reviewed, tested, and edited the resulting text and code as needed and take full responsibility for the content of the published article.

## Materials and Methods

### *C. elegans* strains and maintenance

*C. elegans* strains were maintained on nematode growth medium (NGM) plates seeded with *Escherichia coli* OP50 using standard methods. Unless otherwise indicated, animals were cultured at 20°C. Experiments examining temperature-dependent induction were performed at 25°C, followed, where indicated, by transfer to 20°C for analysis of persistence and inheritance across generations. The CGC1 (https://cgc.umn.edu/strain/CGC1) strain was used as the wild-type reference. All strains used in this study are listed in Supplemental table 1.

### Construction of ZNF-236-prion-domain fusion alleles

Five *znf-236* fusion alleles were generated by CRISPR/Cas9-mediated genome editing in *C. elegans*. All alleles were produced using the same general injection and screening strategy; only the guide RNAs and homologous recombination (HR) templates differed among constructs. The allele names, fusion orientations, inserted domains or tags, guide RNA sequences, HR-template sequences, and genotyping primers are listed in Supplemental table 2 and Supplemental table 3.

#### Guide RNA–Cas9 ribonucleoprotein assembly

For each allele, a target-specific *znf-236* crRNA was ordered from Integrated DNA Technologies (IDT) and annealed with tracrRNA (IDT) to generate the functional guide RNA. A guide RNA targeting the co-conversion marker *dpy-10* was prepared in parallel. The *znf-236* and *dpy-10* guide RNAs were combined with Alt-R™ *S. pyogenes* Cas9 Nuclease V3 (IDT) and incubated at room temperature to form ribonucleoprotein complexes. The guide RNA sequences used for each allele are provided in Supplemental table 2.

#### Homologous recombination templates

HR templates were either generated by PCR amplification from plasmid or synthetic DNA templates or obtained as synthetic double-stranded DNA fragments from Twist Bioscience. Each template contained the intended fusion sequence flanked by short homology arms corresponding to the targeted region of *znf-236*. PCR-amplified templates were generated using Phusion High-Fidelity DNA Polymerase Master Mix (Thermo Fisher Scientific, cat. no. F531L) and purified using the NucleoSpin Gel and PCR Clean-Up Kit (Takara Bio USA, cat. no. 740609). Template source, fusion architecture, homology-arm sequences, amplification primers where applicable, and complete HR-template sequences are provided in Supplemental table 2.

Injected animals were recovered individually, and their F1 progeny were screened for the *dpy-10* co-conversion phenotype (Arribere et al., 2014). Candidate edited animals were isolated and propagated. Precise insertion of each fusion was verified by PCR amplification across the 5′ and 3′ genome–insert junctions, followed by Sanger sequencing of the complete edited region. Verified lines were outcrossed once before use in subsequent experiments.

### RNA-seq sample collection and sequencing

L4-stage [*AZA*^+^] and [*aza*^−^] siblings were identified based on the presence or absence, respectively, of hsp-16.41 reporter expression and collected separately. Two independent experiments were performed, each comprising three biological replicates per state. Each biological replicate consisted of 100 L4-stage animals. Total RNA was extracted using the Direct-zol RNA Miniprep Kit (Zymo Research; catalog no. R2050) according to the manufacturer’s instructions, including DNase treatment. RNA concentration was measured using a Qubit fluorometer. RNA samples were combined with SEQguard Dino Preserve and shipped at room temperature according to Plasmidsaurus sample-submission instructions.

Strand-specific RNA-seq libraries were prepared by Plasmidsaurus using a 3′ poly(A)+ end-counting workflow. Polyadenylated RNA molecules were primed using an oligo-dT primer containing sample barcodes and unique molecular identifiers (UMIs), followed by reverse transcription, second-strand synthesis, tagmentation, indexing, and PCR amplification. Libraries were sequenced on an Illumina NovaSeq X Plus instrument (Illumina; catalog no. 20084804), generating approximately 90-nt single-end cDNA reads. Reads were enriched near the 3′ ends of captured transcripts, with most coverage occurring within approximately the final 400 nt. The six libraries from the first experiment corresponded to Plasmidsaurus sequencing project 5Q64RD (Exp. 1), and the six libraries from the second experiment corresponded to project JNZLY8 (Exp. 2).

### RNA-seq read processing and analysis

All computational analyses were performed using Python v3.11.15 and R v4.4.2. Raw sequencing reads were evaluated using FastQC v0.12.1, and quality-control results were summarized using MultiQC v1.35. Before alignment, FASTQ read headers were standardized by retaining only the first whitespace-delimited field of each read identifier while leaving the nucleotide sequences and corresponding quality scores unchanged.

Approximately 90-nt single-end cDNA reads were aligned using HISAT2 v2.2.2 to the CGC1 draft genome assembly (CGC1_draft_genome_20230804.fa), obtained from the CGC1 reference-genome data repository (https://osf.io/n3fdy/ (Ichikawa et al., 2025). Resulting alignments were coordinate-sorted and indexed using SAMtools v1.22.1. Alignment and sample identities were maintained using the authoritative sample-to-FASTQ mapping, in which induced_1–induced_3 corresponded to [*AZA*^+^] biological replicates and control_1–control_3 corresponded to [*aza*^−^] biological replicates.

#### Gene annotation and read counting

The Liftoff-derived annotation of protein-coding gene models transferred from the C. elegans N2 WS292 reference to the CGC1 genome assembly (N2_WS292.to.CGC1.prot_cod_genes.gtf) was obtained from the CGC1 reference-genome data repository (Ichikawa et al., 2025). Reads overlapping annotated exons were counted using featureCounts v2.1.1 with the settings -F GTF -t exon -g gene_id -s 1. The -s 1 setting was used because the sequencing libraries were forward-stranded. Gene counts were assigned to stable WormBase gene identifiers, and readable gene names were subsequently added from the same Liftoff-derived annotation. Genes lacking an annotated gene name were retained under their WormBase identifiers.

#### Differential-expression analysis

Differential-expression analysis was performed separately for each experiment in R v4.4.2 using DESeq2 v1.46.0 (Love et al., 2014). Raw integer gene counts and sample metadata were used to construct a DESeq2 dataset with the design formula ∼ condition. Genes with zero counts across all six libraries in an experiment were removed; no additional count threshold was applied before model fitting.

The [*aza*^−^] state was specified as the reference condition, and differential expression was calculated using the contrast [*AZA*^+^] versus [*aza*^−^]. Consequently, positive log_2_ fold-change values indicate higher expression in [*AZA*^+^] animals, whereas negative values indicate higher expression in [*aza*^−^] animals. DESeq2-adjusted P values were calculated using i–Hochberg multiple hypothesis correction (additional multiple hypothesis correction tests were…). Variance-stabilizing transformation was performed with blind = FALSE, and principal-component analysis was based on the 500 genes with the greatest variance.

#### Integration of the two RNA-seq experiments

To identify reproducible expression differences, DESeq2 results from the two experiments were joined using stable WormBase gene identifiers. A gene was considered reproducibly differentially expressed if it had a nominal P value < 0.05 in each experiment and showed the same direction of change in both experiments. The two nominal P values were then combined using Fisher’s method, and the resulting combined P values were corrected across all tested genes using Benjamini–Hochberg multiple hypothesis correction. Genes meeting the individual-experiment criteria and having a combined false-discovery rate < 0.05 were designated high-confidence differentially expressed genes. A more stringent effect-size subset was defined as genes that also exhibited an absolute log_2_ fold change ≥ 1, corresponding to at least a twofold change, in both experiments.

Using these criteria, 128 genes were classified as “high-confidence” differentially expressed genes, including 16 with higher expression in [*AZA*^+^] animals and 112 with higher expression in [*aza*^−^] animals. Forty-one of these genes showed at least a twofold change in both experiments.

#### Expression visualization and selection of genes for display

For gene-level expression plots, reads per million (RPM) were calculated separately for each library by dividing the raw count for each gene by the sum of gene-level counts in that library and multiplying by one million. Values were transformed as log_2_(RPM) without adding a pseudocount; measurements with zero counts were therefore treated as missing on the logarithmic scale. The resulting values were imported into GraphPad Prism 10 (GraphPad Software), which was used to generate the gene-level expression plots. Individual points represent biological replicates.

For the gene-expression panel in Figure 4, the 10 high-confidence genes with the largest positive log_2_ fold changes and the 10 with the largest negative log_2_ fold changes in Exp. 1 (5Q64RD) were selected for display. *znf-236* was included separately for comparison. High-confidence genes were defined using the combined two-experiment analysis described above.

MA plots were generated in Python using Matplotlib and show the DESeq2 log_2_ fold change as a function of log_2_(baseMean+1), where baseMean is the mean DESeq2-normalized count across all samples in the corresponding experiment. Genes were classified as differentially expressed using the adjusted-P-value and fold-change thresholds indicated in the figure legend. Selected high-confidence genes, heat-shock protein genes, and *znf-236* were annotated on the plots.

